# SMARCA4 supports the oncogenic landscape of *KRAS*-driven lung tumors

**DOI:** 10.1101/2020.04.18.043927

**Authors:** Shivani Malik, Masaya Oshima, Nilotpal Roy, Swati Kaushik, Ora Kuvshinova, Wei Wu, John E. Greer, Shon Green, Martin McMahon, Kuang-Yu Jen, Matthias Hebrok, Sourav Bandyopadhyay, Diana C. Hargreaves, Eric A. Collisson

**Affiliations:** Division of Hematology and Oncology, Department of Medicine, University of California, San Francisco, USA; Molecular and Cell Biology Laboratory, Salk Institute for Biological Studies. La Jolla, USA; Diabetes Center, Department of Medicine, University of California, San Francisco, USA; UCSF Helen Diller Family Comprehensive Cancer Center, San Francisco, USA; Huntsman Cancer Institute, 2000 Circle of Hope, Salt Lake City, USA; Department of Pathology, University of California, San Francisco, USA

## Abstract

Cancer resequencing studies identify recurrent mutations in the switch/sucrose non-fermentable (SWI/SNF) complex at an unexpectedly high frequency across many cancer types. Some SWI/SNF mutations appear to be loss-of-function events, implying that the intact SWI/SNF complex is tumor suppressive. We examined the distribution and function of *SMARCA4* mutations, the most frequently mutated SWI/SNF complex gene in lung adenocarcinoma, using human cancers, cell lines and mouse model systems. We found that lung adenocarcinomas harboring activated oncogenes have fewer deleterious mutations in *SMARCA4* and express higher levels of the mRNA than cancers without activated oncogenes, indicating distinct dependencies on *SMARCA4* in these two settings. Surprisingly, intact *Smarca4* promoted the growth and tumorgenicity of *Kras*^*G12D*^-driven mouse lung tumors and human cells. Mechanistically, we found that *Smarca4* supports the oncogenic transcriptional/signaling landscape of *Kras*^*G12D*^-driven mouse lung cancer. This dependency on the chromatin maintenance machinery in established cancer cells support treatments directed towards pathogenic SWI/SNF complexes in lung adenocarcinoma and other malignancies.

## Introduction

Lung adenocarcinoma (ADC) is, by far, the leading cause of cancer death. The American Cancer Society estimates that about 156,000 people died of lung cancer in the United States in 2017, claiming more lives than prostate, breast and colon cancer combined. Next generation sequencing has enabled comprehensive analysis of genetic alterations in lung ADCs and has uncovered alterations in oncogenic driver genes in the RTK-RAS-RAF pathway. These activating, gain of function mutations (such as *KRAS, EGFR, BRAF, ERBB2* mutations and *ALK, NTRK* and *ROS1* fusions) can be identified in up to 75% of lung ADCs, categorizing them as the “oncogene positive” subset of lung ADCs. The remaining 25% of lung ADCs without any detectable alteration in known oncogenic drivers are termed the “oncogene negative” subset of lung ADCs.

Small molecule inhibitors have been used to successfully target many of these oncogenic drivers and such targeted therapies have shown to be more effective and less toxic than standard chemotherapy. For example, erlotinib, afatinib, gefitinib and osimertinib are all FDA-approved for patients whose tumors harbor *EGFR* activating mutations. Similarly, *ALK*- and *ROS1*-fusion driven lung ADCs respond well to targeted inhibitors. However, developing therapeutics for oncogene-negative lung ADC patients is a huge unmet challenge. In order to study the oncogene negative subset of lung ADC, we looked for genes whose mutation rates were higher in oncogene-negative lung ADCs compared to oncogene-positive ADCs. We find *SMARCA4* mutations are significantly more frequent in the oncogene negative compared to oncogene positive lung ADCs. We further find that KRAS-addicted lung ADCs, compared to *KRAS* independent lung ADCs, harbor relatively lower rates of *SMARCA4* mutation.

SWI/SNF complexes form upon assembly of 12-15 subunits around one of the two ATPase subunits and are 2 mega Daltons fully assembled. The two ATPase subunits are encoded by *SMARCA4* (protein-BRG1) or *SMARCA2* (protein-BRM). The DNA-dependent, ATPase activity of BRG1 or BRM can either activate or repress mRNA transcriptional programs by controlling chromatin accessibility through nucleosome positioning and ejection and in doing so, impact cellular growth and development^1^.

Intact *SMARCA4* cooperates with oncogenic *KRAS* to promote pancreatic intraepithelial neoplasia and pancreatic ductal adenocarcinoma as well as melanoma^2–4^. This finding led us to hypothesize that the SWI/SNF complex in general and *SMARCA4* specifically may paradoxically support, rather than constrain, *KRAS-*driven lung adenocarcinoma. We tested this hypothesis by examining *Kras*-driven mouse lung cancers in the conditional presence or absence of *Smarca4*. We show that *Smarca4* cooperates with oncogenic *Kras* to promote growth and malignancy of *Kras*^*G12D*^-driven lung tumors, using both synchronous and temporally separated *Kras* activation and *Smarca4* inactivation. In an orthologous system, we find that depletion of SMARCA4 reduces transformative potential of KRAS^G12D^-transformed human airway cells. These results together show that unlike traditional tumor suppressor genes (TSGs), loss of *SMARCA4* antagonizes KRAS signaling explaining the positive selection of *SMARCA4* mutations in oncogene negative subset of lung ADCs.

Over the last decade, both gain-of-function and loss-of-function alterations in SWI/SNF complex have been described in cancer. For example, in synovial sarcoma fusion of 78 amino acids of SSX protein with SS18 subunit of SW/SNF complex targets the SWI/SNF complex to loci such as *SOX2*, leading to increased cell proliferation^5^. In rhabdoid tumors on the other hand, biallelic inactivation of *BAF47* destabilizes the SWI/SNF complex leading to loss of enhancer regulation^6,7^. However, functional and mechanistic contributions that SWI/SNF mutations make to the genesis and maintenance of common solid tumors has been more difficult to interpret. For example, while considered to be a tumor suppressor, *Smarca4* is required for Wnt-driven adenoma formation in the epithelium of small intestine^8^, but is recurrently mutated in small cell carcinoma of the ovary^9–11^. Similarly, a SMARCA4-driven BAF complex is required for proliferation of leukemia cells and its manipulation has been proposed to be a potential therapeutic strategy^12^. Recent studies have also highlighted context-specific and dosage-dependent roles of SWI/SNF subunits SMARCA4^2,3^ and ARID1A (BAF250)^13^ in pancreatic and hepatocellular carcinomas respectively implying to a more nuanced regulation of the cancer-specific properties of SWI/SNF complexes, possibly based on stage of the disease.

We find that loss of *SMARCA4* dampens KRAS signaling, impairs cellular proliferation and decreases malignant progression of KRAS-initiated lung tumors. This study identifies SMARCA4 as a potential specific vulnerability in KRAS-driven lung cancer. Further, the signaling/transcriptional landscape engendered by the loss of *Smarca4* may be uniquely operational in the oncogene negative subset of lung ADCs, a currently poorly understood and ineffectively treated subset of this deadly disease.

## Results

### *SMARCA4* mutations are enriched in the subset of lung adenocarcinoma patients

To search for pathways important in lung adenocarcinomas without identified oncogene alterations, we compared mutation rates of significantly mutated genes between tumors with (n=295) and without (n=261) activated oncogene (Figure 1A, left). We found that *SMARCA4* was mutated more much more frequently in oncogene negative samples than in oncogene positive cases (5.4% vs 12.1%, p = 0.01) (Fig. 1A, right), indicating a potential requirement of intact *SMARCA4* in oncogene-driven lung ADC. Singh et al.^14^ previously defined a *KRAS* dependent gene signature in lung ADC. To assess the integration between KRAS dependency and SMARCA4 mutation, we next examined *SMARCA4* mutational frequency in lung ADCs bearing *KRAS* mutation and ranked by a *KRAS*-high/dependent gene signature (Figure 1B, left). We found *SMARCA4* mutations less frequently in KRAS-dependent lung ADCs compared to KRAS-independent (KRAS low signature) lung ADC patients (Fig. 1B right). Notably, missense S*MARCA4* mutations, which retained *SMARCA4* expression, were enriched in the KRAS-dependent lung ADC cohort (32.5% vs 18.6%, p= 0.0051). On the other hand, deleterious mutations (nonsense, frameshift and splice site mutations), which lead to loss of expression, were enriched in the KRAS-independent lung ADCs (9.3% vs 39.5%, p =0.0051, Fig. 1B right).

**Figure 1.**
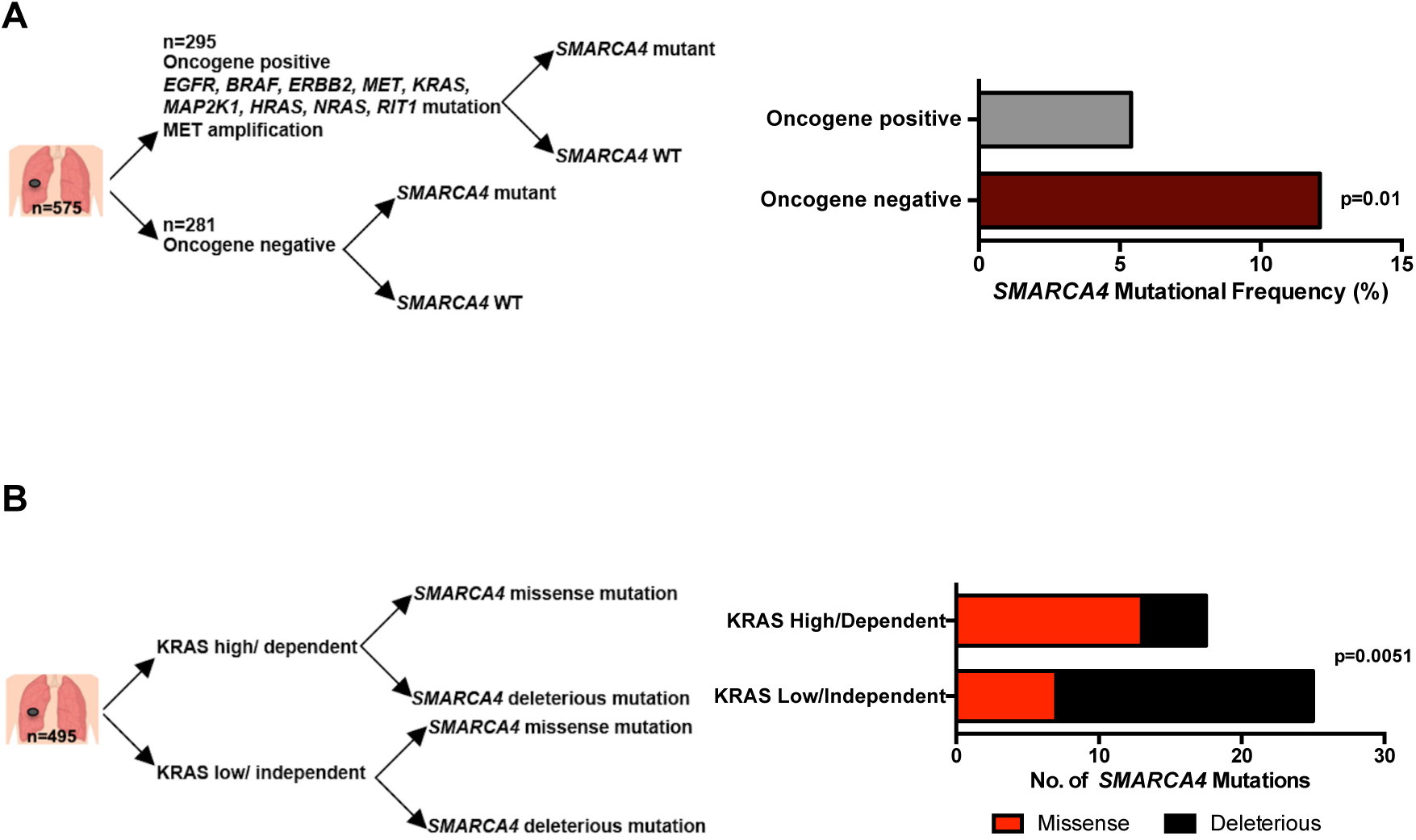
SMARCA4 mutations are enriched in oncogene negative subset of lung adenocarcinoma (ADC) patients. (A) Left-Schematic of analysis. ADCs were divided into oncopositive or onconegative based on presence of oncogenic mutations (see Methods). Right-*SMARCA4* mutational frequency in onconegative group of ADCs is significantly (Fisher exact test, p=0.01) higher than oncopositve group. (B) Left-Schematic of analysis-lung ADCs were divided into KRAS dependent/high or independent/low groups based on KRAS transcriptional signature. Right-significantly higher number of missense and lower number of deleterious *SMARCA4* mutations observed in KRAS high ADCs compared to KRAS low ADCs (Fisher exact test, p value=0.005).

### SMARCA4 loss impedes malignancy of KRAS^G12D^-driven mouse lung tumors

This observation of infrequent loss of *SMARCA4* in oncogenic KRAS-driven lung ADCs led us to hypothesize that SMARCA4 may have tumor supporting roles in oncogenic KRAS-dependent lung tumor development. We tested this hypothesis *in vivo* by generating a GEMM carrying *Lox-stop-Lox Kras*^*G12D*^ (*Kras*^*LSL_G12D* ref. 15^) and homozygous, floxed alleles of *Smarca4* (*Smarca4*^*f/f* ref. 16^) (Fig. 2A). Cre recombinase (delivered through adenovirus expressing Cre; Adeno-Cre) excises the Lox-stop-Lox cassette in the *Kras*^*LSL_G12D*^ allele and induces expression of *Kras*^*G12D*^ from the endogenous promoter. Exons 2 and 3 of *Smarca4* are concurrently excised, eliminating its expression in Adeno-Cre infected lung cells (Fig. 1C). We tested if SMARCA4 plays any role in RAS signaling by assessing RAS effectors signaling in *Kras*^*LSL_G12D*^; *Smarca4*^*+/+*^ *and Kras*^*LSL_G12D*^; *Smarca4*^*f/f*^ mice at 2 different time points (3 and 6 months) after Adeno-Cre infection (Fig. 2A).

**Figure 2.**
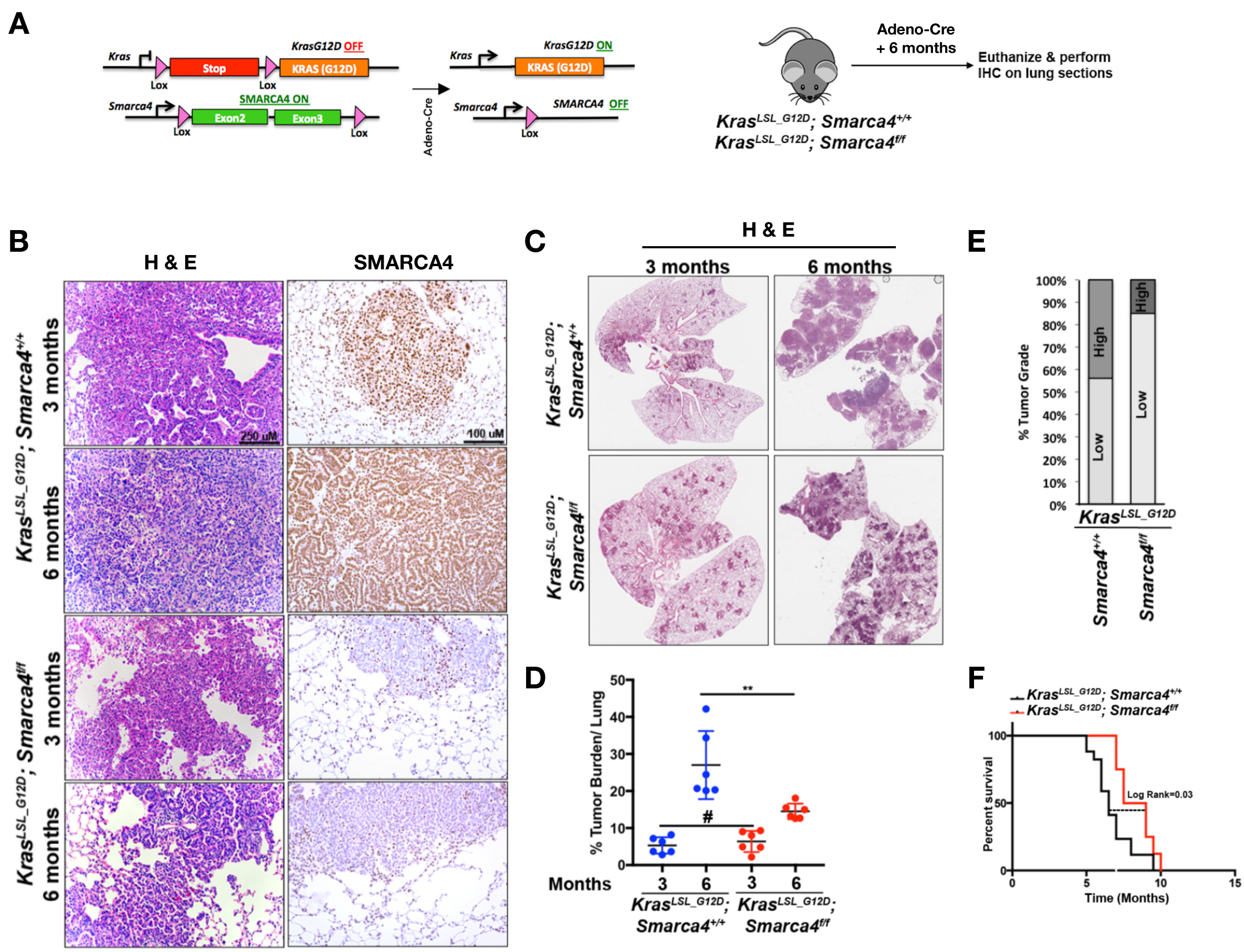
SMARCA4 loss delays KRASG12D driven mouse lung tumor progression. (A) Left-Mouse alleles employed for generating genetically engineered mouse model (GEMM). Right-Schematic of mouse experiment performed for obtaining tumor. (B) Representative H&E and Immunohistochemistry (IHC) of SMARCA4 of indicated genotypes from *Kras*^*LSL_G12D*^; *Smarca4*^*+/+*^ or *Kras*^*LSL_G12D*^;*Smarca4*^*f/f*^ mice euthanized 3 months or 6 months post Adeno-Cre infection to assess lung tumor burden (C) Representative Hematoxylin and Eosin (H&E)-stained lung sections at indicated time points. (D) Quantification of lung tumor burden (tumor area/total area of lung) of mice at various time points post Adeno-Cre infection. (n=6 for each time point). Error bars indicate s.e.m. Significance was determined using Student’s two-tailed t-test. (E) Quantification of grades of *Kras*^*LSL_G12D*^; *Smarca4*^*+/+*^ (n=5) and *Kras*^*LSL_G12D*^; *Smarca4*^*f/f*^ (n=6) tumors. (F) Kaplan-Meir curve of the indicated genotypes. Significance was calculated using log rank test. *Kras*^*LSL_G12D*^; *Brg1*^*+/+*^ (n=17), *Kras*^*LSL_G12D*^; *Brg1*^*f/f*^ (n=8). All error bars represent s.e.m.

We examined tumors from *Kras*^*LSL_G12D*^; *Smarca4*^*+/+*^ and *Kras*^*LSL_G12D*^; *Smarca4*^*f/f*^ mice at 3 months and 6 months post Adeno-Cre infection in order to assess the role of *Smarca4* in lung tumor genesis. Immunohistological staining of *Kras*^*LSL_G12D*^; *Smarca4*^*f/f*^ tumors showed about 85% of tumors had no staining of SMARCA4 upon Adeno-Cre infection (Fig. 2B). *Kras*^*LSL_G12D*^; *Smarca4*^*+/+*^ and *Kras*^*LSL_G12D*^; *Smarca4*^*f/f*^ mice displayed similar tumor burden 3 months post Adeno-Cre infection (Figure 2C-D). Interestingly, tumor burden in *Kras*^*LSL_G12D*^; *Smarca4*^*+/+*^ mice was greater 6 months post infection, compared to *Kras*^*LSL_G12D*^; *Smarca4*^*f/f*^ animals (Fig. 2C-D). 45% of the *Kras*^*LSL_G12D*^; *Smarca4*^*+/+*^ tumors were high-grade papillary adenomas whereas only 15% of *Kras*^*LSL_G12D*^; *Smarca4*^*f/f*^ tumors were high-grade (Fig. 2E). High grade tumors in *Kras*^*LSL_G12D*^;*Smarca4*^*+/+*^ mice correlated with their relatively shorter overall survival compared to *Kras*^*LSL_G12D*^; *Smarca4*^*f/f*^ mice (Fig. 2F).

Previous studies have correlated transcription factor expression patterns to tumor grade in oncogenic *Kras*^*G12D*^-driven lung cancer^17^. For example, high levels expression of Ttf1 (encoded by *Nkx2-1*) and Foxa2 correlated with low tumor grade of oncogenic KRAS-driven lung tumors while Hmga2 was expressed exclusively in high-grade tumors^17^. The expression of these factors thus can be used as markers of progression from benign adenomas to malignant forms. Immunohistological staining showed decrease TTF1 and FOXA2 expression between 3 months and 6 months after Adeno-Cre infection in *Kras*^*LSL_G12D*^; *Smarca4*^*+/+*^ lung tumors (Fig. 3A) correlated with a suppressive role of these factors in lung tumor progression. In contrast, strong Ttf1 and Foxa2 expression were maintained in *Kras*^*LSL_G12D*^; *Smarca4*^*f/f*^ tumors after 6 months of Adeno-Cre infection (Fig. 3A). The expression of Hmg2a was detected in 2/6 *Kras*^*LSL_G12D*^; *Smarca4*^*+/+*^ tumors at 6 months post Adeno-Cre infection, specifically in high grade tumors) but absent from *Kras*^*LSL_G12D*^; *Smarca4*^*+/+*^ tumors at 3 months and also not found in *Kras*^*LSL_G12D*^; *Smarca4*^*f/f*^ tumors at 6 months post Adeno-Cre (Fig. 3A). Additionally, immunohistochemical staining revealed reduced levels of phosphorylated AKT (p-AKT) and phosphorylated ERK1/2 (p-ERK1/2) in *Kras*^*LSL_G12D*^; *Smarca4*^*f/f*^ tumors compared to *Kras*^*LSL_G12D*^; *Smarca4*^*+/+*^ lung tumors (Fig. 3B), consistent with a lower tumor grade.

**Figure 3.**
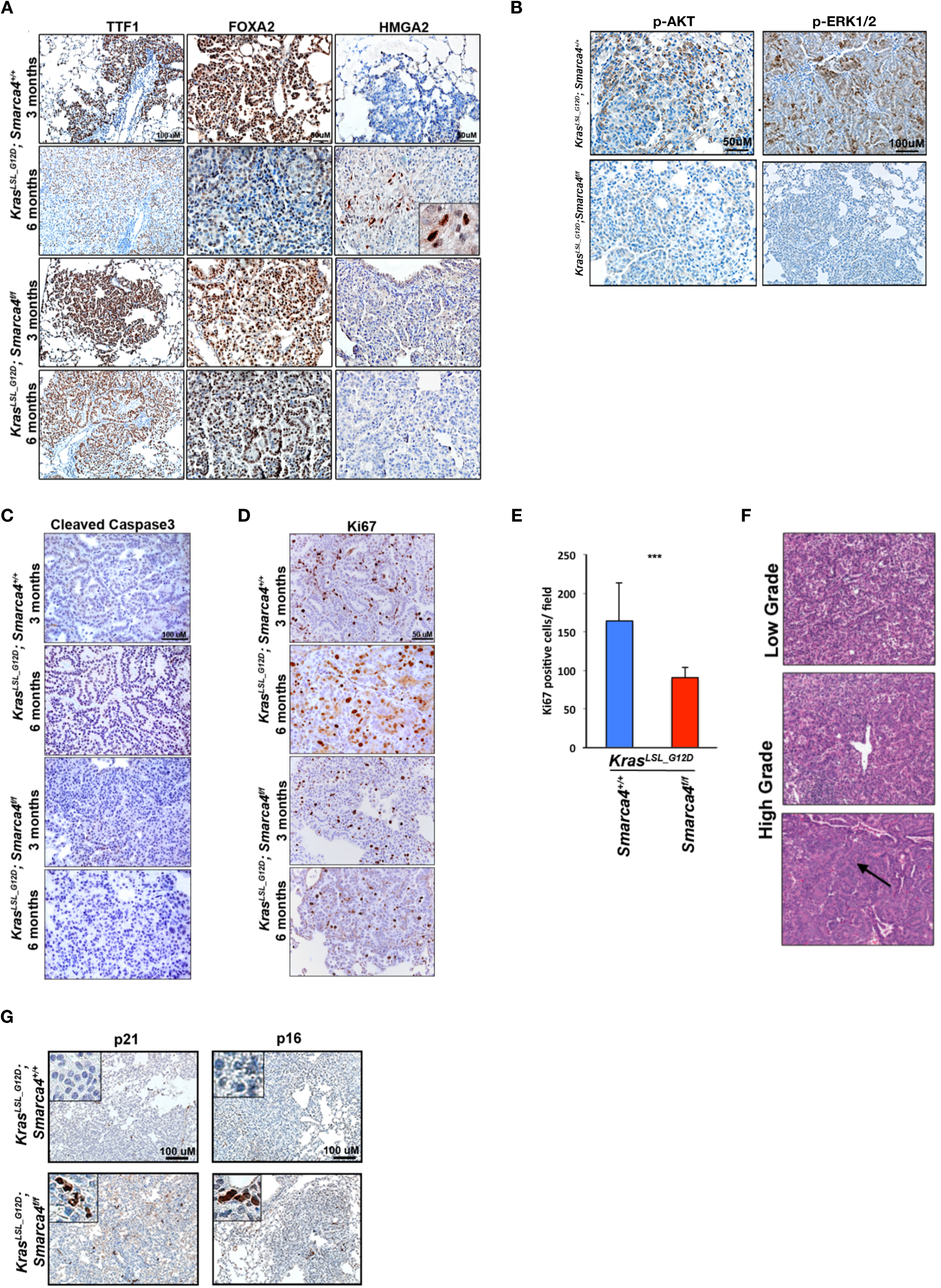
SMARCA4 loss impedes malignancy of KRASG12D-driven mouse lung tumors tumors. (A) IHC staining of TTF1, FOXA2 and HMGA2 in lung tumors of indicated genotypes at indicated time points post Adeno-Cre infection. (B) p-AKT and p-ERK1/2 staining in sections of mouse lung tumors of indicated genotypes 6 months post Adeno Cre. (C-D) Representative IHC of Cleaved Caspase 3 and Ki67 of indicated genotypes at indicated time points. (E) Quantification of Ki67 positive cells in lung tumors of indicated genotypes (n=12 random fields). Significance calculated by paired two-tailed Student’s *t*-test between the respective genotypes is indicated. #-not significant, **<0.01. (F) Grading scheme followed. Top: Papillary adenoma showing cellular proliferation with papillary architecture and small, bland nuclei (low-grade). Middle: A small focus of high-grade, pleomorphic nuclei with prominent nucleoli and scattered intranuclear inclusions; this focus likely represents malignant transformation within a papillary adenoma. Bottom: Papillary adenocarcinoma showing widespread high-grade, pleomorphic nuclei with prominent nucleoli and readily identifiable mitotic figures (arrow). (G) IHC staining of senescence markers p21^Waf1/Cip1^ and p16^INK4a^ in lung tumors of mice of indicated genotypes 3 months post adeno-Cre infection.

As *Smarca4* loss decreased activation of AKT and MAPK, two major pathways in cell survival, proliferation and tumorigenesis, we looked whether loss of SMARCA4 induced cell death or impede proliferation. We did not observe apoptosis as measured by cleaved caspase 3 staining in lung sections from either genotype (Fig. 3C). On the other hand, *Kras*^*LSL_G12D*^; *Smarca4*^*f/f*^ tumors were less proliferative than *Kras*^*LSL_G12D*^; *Smarca4*^*+/+*^ tumors as indicated by a lower Ki67 index (Fig. 3D-E). Additionally, the majority of *Kras*^*LSL_G12D*^; *Smarca4*^*+/+*^ mice showed features of malignant transformation such as readily identifiable mitotic figures whereas *Kras*^*LSL_G12D*^; *Smarca4*^*f/f*^ tumors mostly depicted small and bland nuclei (Fig. 3F). Remarkably, 0 out of 6 *Kras*^*LSL_G12D*^; *Smarca4*^*f/f*^ mice showed malignant features such mitotic figures (Fig. 3F). To strengthen our observation, we also looked to markers of cell cycle arrest p21^Waf1/Cip1^ and p16^INK4a^. Consistent with slower proliferation in the SMARCA4 knockout tumors, we observed high levels of senescence markers, p16^INK4a^ and p21^Waf1/Cip1^ in the *Kras*^*LSL_G12D*^; *Smarca4*^*f/f*^ tumors compared to *Smarca4* intact tumors (Fig. 3G). These findings suggest that SMARCA4 accelerates the progression from precursor lesions to a malignant form.

### *Smarca4* is selectively retained in *Kras*^*G12D*^-transfomed lung epithelial cells during lung tumor development

Our data suggest a role of SMARCA4 supporting oncogenic KRAS during tumor progression. We next asked if *Smarca4* was indispensable for the maintenance of lung tumor cells transformed by *Kras*^*G12D*^ by using *Kras*^*FrtStopFrt_G12D*^;*Spc*^*CreERT2*^;*Smarca4*^*flox/flox*^ mice (Fig. 4A).

**Figure 4.**
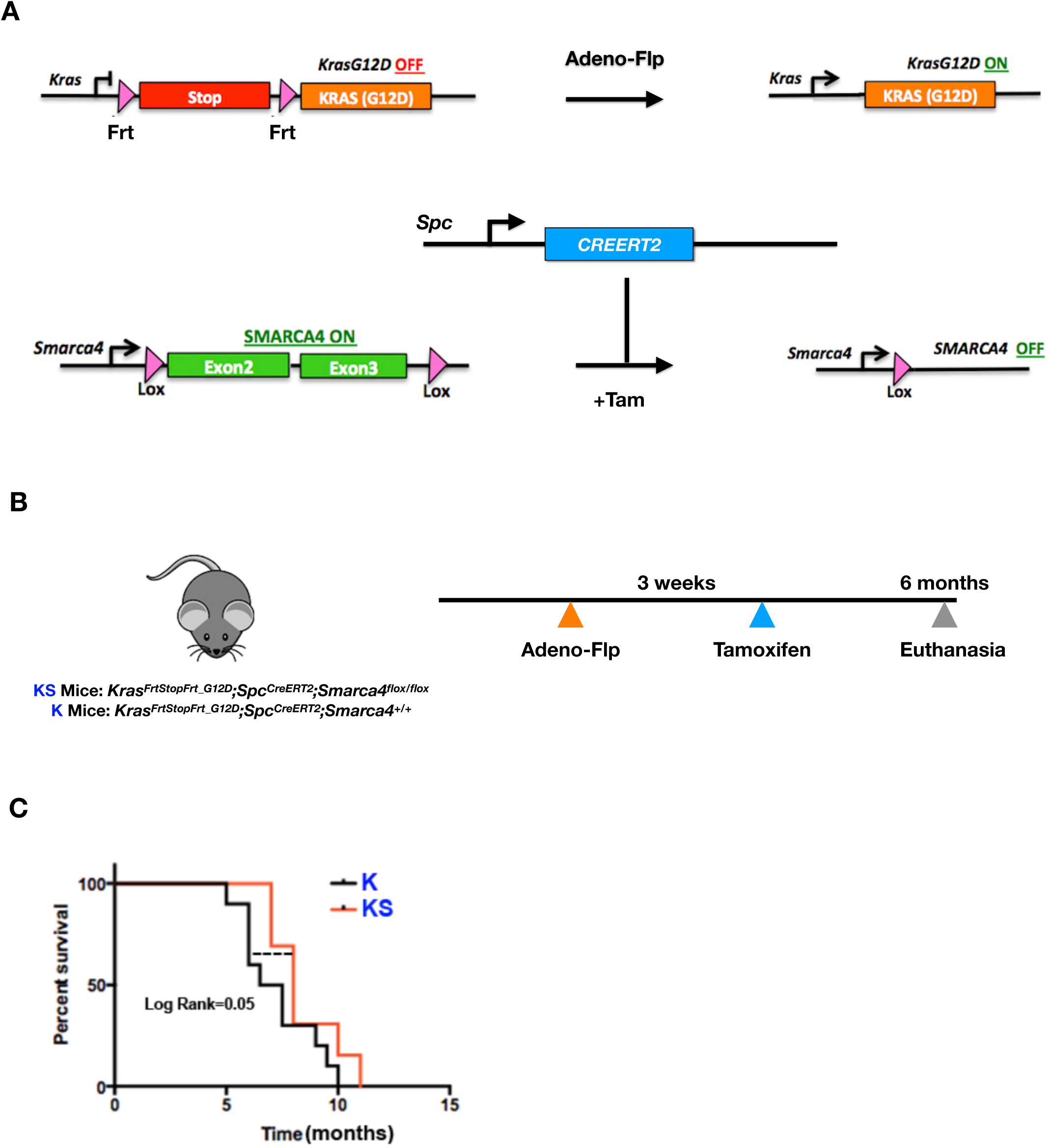
*Smarca4* is selectively retained in KRAS^G12D^-transfomed lung epithelial cells during lung tumor development. (A) Mouse alleles employed (B) experimental strategy (C) Kaplan-Meir curve of the indicated mice.

In these mice, expression of oncogenic *Kras*^*G12D*^ can be induced in lung using Adeno viruses encoding Flippase. Homozygous *Smarca4* knockout was induced after 4-hydoxytamoxifen (Tam)^18^ induced stabilization of CreERT2 in *Spc* (Surfactant Protein C) expressing lung epithelial cells. *Spc* is expressed uniformly by type II lung pneumocytes, the major cell of origin of lung adenocarcinoma^19^ and therefore appropriate for our experiments.

We compared survival of *Kras*^*FrtStopFrt_G12D*^;*Spc*^*CreERT2*^;*Smarca4*^*+/+*^ mice (hereafter K) to *Kras*^*FSF_G12D*^;*Spc*^*CreERT2*^;*Smarca4*^*flox/flox*^ mice (hereafter KS) to test if loss of *Smarca4* impacted survival of mice bearing *Kras*^*G12D*^-tumors where *Smarca4* knockout was induced 3 weeks post Adeno-Flp infection (Fig. 4B). We observed modest yet significant increase in lifespan of KS compared to K mice (Fig. 4C). This difference is likely an underestimate, as the efficiency of *Smarca4* deletion in the *Spc*^*CreERT2*^ animals was lower of what could be achieved with the *Smarca4*^*flox/flox*^ allele (Supplementary Figure 1). Indeed, biallelic recombination in tumors from KS mice was ∼20% and retained *Smarca4* expression (Supplementary Figure 1). This likely contributes to the modest survival advantage of KS mice compared to K (Fig. 4C).

Together, our findings from temporal GEMM indicate that SMARCA4 supports the oncogenic landscape of KRAS transformed cells.

### SMARCA4 supports Kras^G12D^ signaling in mouse lung tumors

We isolated lung tumors from end stage (body conditioning score=2) *Kras*^*LSL_G12D*^; *Smarca4*^*f/f*^ mice and derived cells from these tumors to study how SMARCA4 mechanistically regulated lung tumorigenesis. We re-expressed *SMARCA4* in these mouse lung tumor cells to generate isogenic cell lines differing only in expression of *SMARCA4* (Fig. 5A). The cell line derived was tested for recombined *Kras* allele (Fig. 5B) and loss of SMARCA4 protein (Fig. 5C). Infection of *Kras*^*LSL_G12D*^; *Smarca4*^*f/f*^ tumor cell line with *SMARCA4* cDNA led to high expression of SMARCA4 protein as assessed by western blot (Fig. 5C). Since SMARCA4 regulates nucleosome occupancy and affects local mRNA transcription^20^, we performed RNA sequencing analysis of control and *SMARCA4* overexpressing cells derived from *Kras*^*LSL_G12D*^;*Smarca4*^*f/f*^ mice lung tumor. Consistent with its role in regulating transcription, overexpression of SMARCA4 led to differential expression of 446 genes (≥1.5 fold change) compared to control cells (Fig. 5D). We found enrichment of gene set up regulated by oncogenic *Kras* (KRAS.600_UP.V1_UP) in the SMARCA4 re-constituted line (Fig. 5E) such as *Inhba*. Gene-set down regulated upon expression of oncogenic KRAS (SWEET_LUNG_CANCER_KRAS_DN) such as *Smad6* was found to be also down regulated in SMARCA4 expressing cell line (Fig. 5E). Expression of ribosomal genes (KEGG_RIBOSOME) was up regulated and protein translation pathway genes (REACTOME_TRANSLATION) were significantly enriched in the *SMARCA4* overexpressing cells, suggesting hyper-activation of PI3’K-AKT pathway (Fig. 5E).

**Figure 5.**
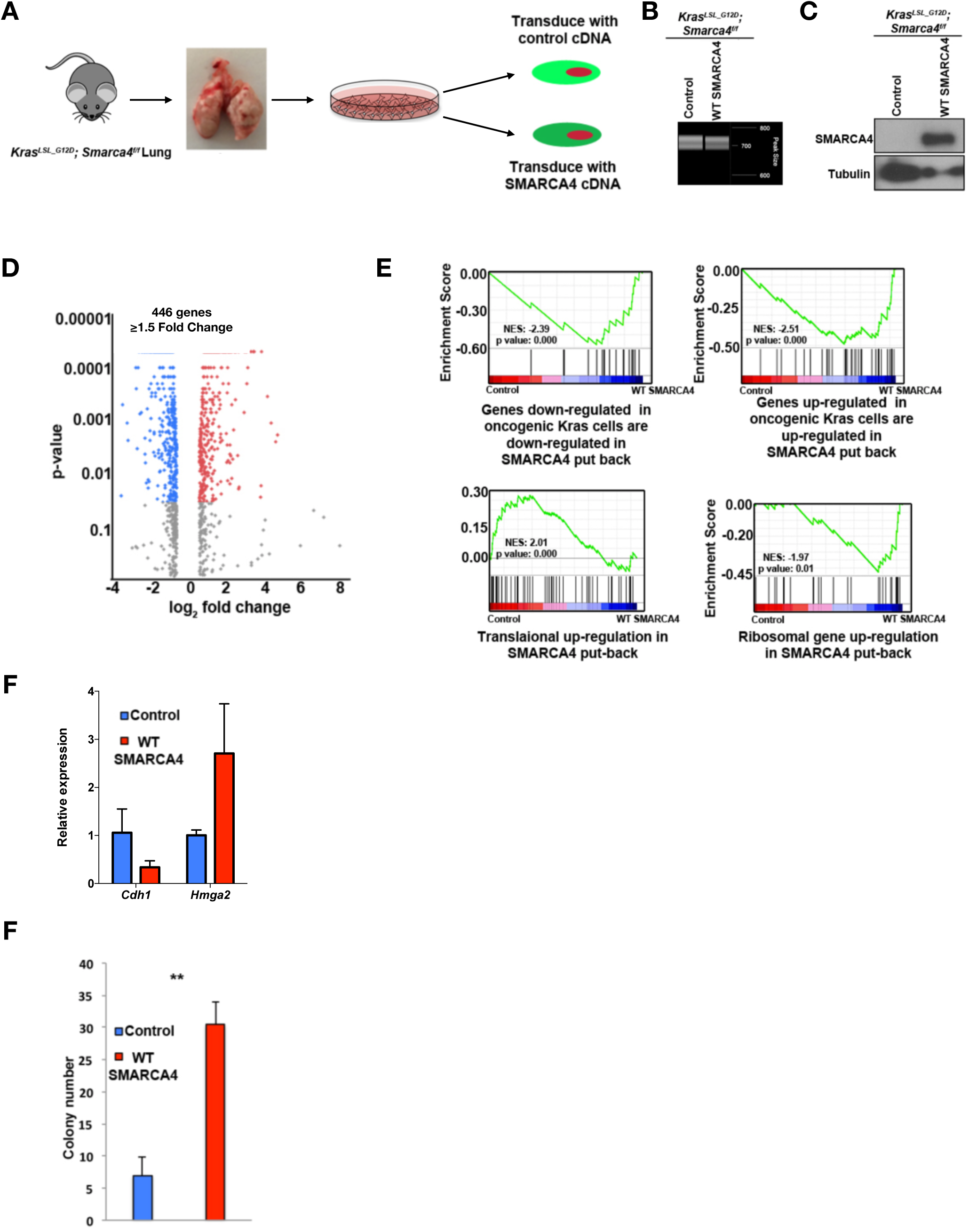
SMARCA4 co-operates with KRAS to maintain active RAS signaling and associated EMT. (A) Schematic of cell line generation from *Kras*^*LSL_G12D*^; *Smarca4*^*f/f*^ tumor. (B) PCR analysis of *Kras*^*G12D*^ allele in cells isolated from lung tumors of *Kras*^*LSL_G12D*^; *Smarca4*^*f/f*^ mice show recombination of the lox sites flanking the stop cassette. (C) Western blot showing ectopic expression of SMARCA4 in SMARCA4 null *Kras*^*LSL_G12D*^; *Smarca4*^*f/f*^ cells. (D) Volcano plot depicting differentially expressed genes between the control (*Smarca4* deficient) and SMARCA4 re-expressing *Kras*^*LSL_G12D*^; *Smarca4*^*f/f*^ cell lines (WT SMARCA4). (E) GSEA of all the genes altered by at least -/+ log 0.6 fold in the SMARCA4 re-expressing *Kras*^*LSL_G12D*^; *Smarca4*^*f/f*^ cell lines compared to control cells. Enrichment of genes up regulated by oncogenic KRAS (KRAS.600_UP.V1._UP) and down regulated by oncogenic Kras (SWEET_LUNG_CANCER_KRAS_DN) in WT SMARCA4 cells. **<0.01. (F) Re-expression of SMARCA4 in *Kras*^*LSL_G12D*^; *Smarca4*^*f/f*^ mouse tumor cells (WT SMARCA4) increases anchorage independent growth on soft agar. Significance calculated using Student’s t-test.

Consistent with immunological staining, *Cdkn1a* (encoding p21) and *Nkx2-1* were downregulated whereas *Hmga2* was upregulated after SMARCA4 overexpression. The epithelial to mesenchymal transition (EMT) pathway was among the most significantly enriched gene sets in the SMARCA4 overexpressing *Kras*^*LSL_G12D*^; *Smarca4*^*f/f*^ mouse cell line (p=10^−22^, hypergeometric test). The epithelial gene *Cdh1* was downregulated whereas mesenchymal gene *Acta2* was upregulated in *Kras*^*LSL_G12D*^; *Smarca4*^*f/f*^ tumor cell line with *SMARCA4* cDNA. These changes suggest that the cDNA reconstitution supports a more malignant expression program. Therefore, we studied whether SMARCA4 rescue induces higher growth rate than in control-Smarca4 knocked-out cells. Overexpression of *SMARCA4* into mouse derived *Kras*^*LSL_G12D*^; *Smarca4*^*f/f*^ cells increased anchorage independent growth of the reconstituted cell line compared to *Kras*^*LSL_G12D*^; *Smarca4*^*f/f*^ control cell line (Fig. 5F), a result consistent with our *in vivo* data (Fig. 2-3).

As a whole, the enrichment of oncogenic KRAS-associated ribosomal and translational gene signatures in SMARCA4 expressing cells strongly suggests that SMARCA4 supports oncogenic KRAS signaling and this may be one of the manner by which SMARCA4 promotes tumorigenicity in mouse models.

### SMARCA4 promotes transformative properties of human lung cancer cells

Given, the growth supportive properties of SMARCA4 in mouse, we next asked if SMARCA4 supported growth of human lung cancer cells. We overexpressed WT *SMARCA4* in human lung adenocarcinoma line, A549 known to harbor a homozygous *SMARCA4* mutation^21^. Overexpression of WT *SMARCA4* in A549 resulted in its robust expression and reconstitution of SMARCA4 containing SWI/SNF complex as tested by IP-western (6A), IP-Mass spectrometry (6B) and glycerol sedimentation assay (Fig 6C).

**Figure 6.**
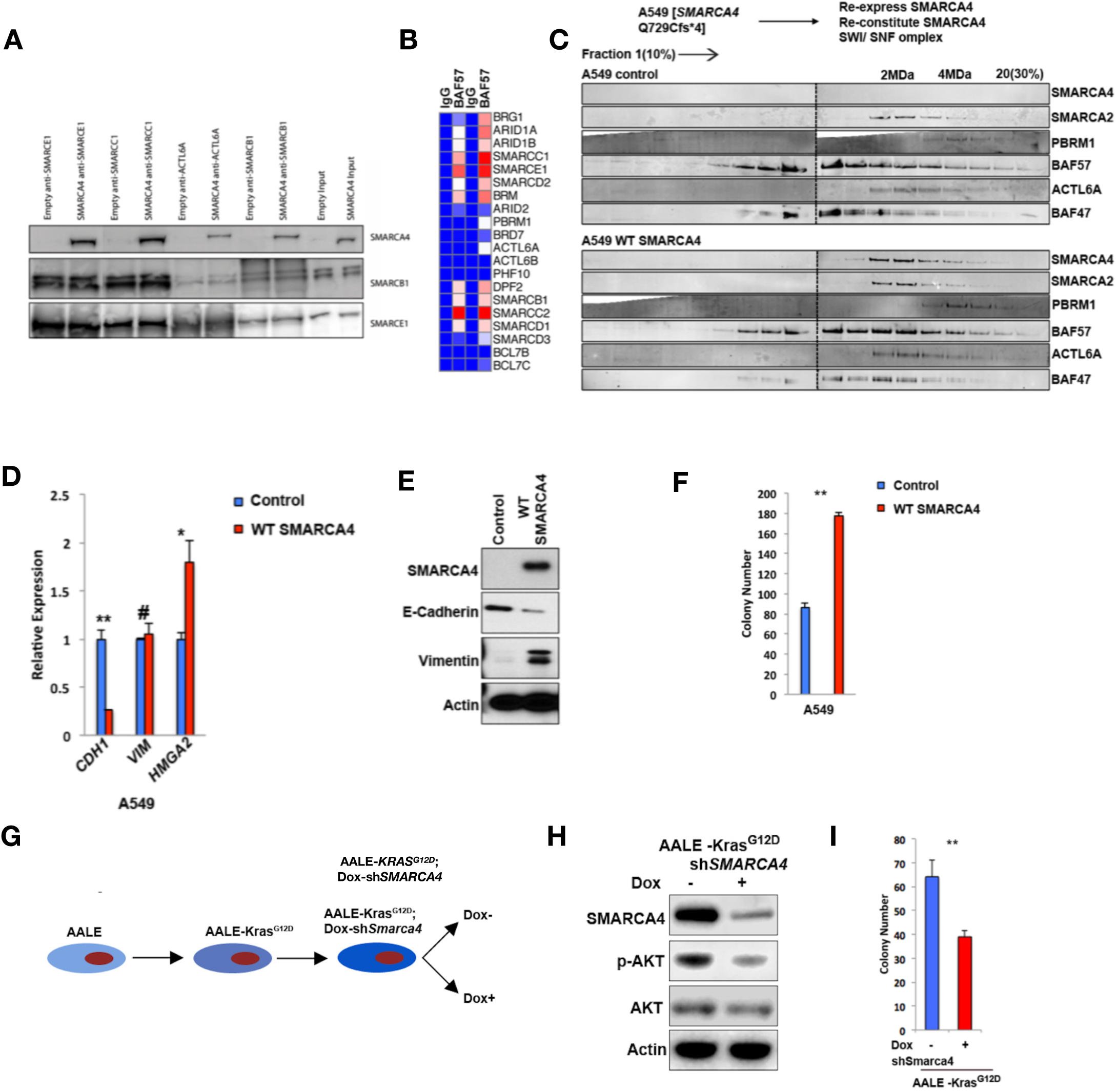
SMARCA4 promotes transformative properties of human lung cells. (A) Immuno-precipitation against the core SWI/SNF subunit, BAF57 was performed followed by western blot (B) Density map showing spectral count of SWI/SNF peptides detected in mass spectrometric analysis of control and SMARCA4 reconstituted A549 extracts. Mass spectrometry was performed following immunoprecipitation of SWI/SNF complex using BAF57 antibody. (C) Western blots of fractions 1-20 of the glycerol gradient sedimentation analysis assessing the migration of BRM-/BRG1-containing BAF complexes at 2MDa and PBAF complexes at 4MDa in A549 control or SMARCA4 reconstituted A549 cells. (D) qPCR (E) Western blot assessing expression of EMT genes in A549 control or SMARCA4 reconstituted A549 cells. (F) Ectopic expression of *SMARCA4* in BRG1 deficient human lung tumor line, A549 augments colony formation in soft agar. (G) Experimental scheme of doxycycline induced SMARCA4 knockdown in AALE-KRAS^G12D^ cells (H) Western blot showing decreased p-AKT upon depletion of SMARCA4 (dox +) in AALE-Kras^G12D^ cells. (I) Doxycycline inducible depletion of *SMARCA4* in AALE-Kras^G12D^ cells reduced colony formation in soft agar. *<0.05, **<0.01.

We compared the expression of *HMGA2, CDH1* and *VIM* in A549 cells expressing control vector or *SMARCA4* at mRNA and protein levels. *HMGA2 expression* was increased in the SMARCA4 expressing A549 cells (Fig. 6D). EMT markers were also affected: epithelial marker *CDH1* mRNA and protein were dramatically decreased in SMARCA4 re-constituted cells (Fig. 6 D-E) whereas mesenchymal VIM protein levels were markedly increased in cells reconstituted with SMARCA4 (Fig. 6E). Consistent with our mice cells findings (Fig. 5F), WT *SMARCA4* expressing A549 cells have increased anchorage independent growth compared to control cell line harboring *SMARCA4* mutation (Fig. 6F).

To support our observation, we utilized immortalized human airway cells (AALE) to examine if SMARCA4 plays similar supportive role in KRAS signaling in human cells. We overexpressed *KRAS*^*G12D*^ in AALE cells and then further engineered them to express a doxycycline-inducible shRNA directed against *SMARCA4* (Fig. 6G). Addition of doxycycline (dox) reduced SMARCA4 protein level by ∼80% compared to control (no dox) as measured by western blot (Fig. 6H). We found p-AKT levels reduced by 60% in SMARCA4 depleted KRAS^G12D^-AALE cells compared to control KRAS^G12D^-AALE cells (Fig. 6H). Finally, we assayed anchorage-independence in KRAS^G12D^-transformed AALE cells expressing tetracycline-inducible shRNA against *SMARCA4* to test tumor supportive role of SMARCA4 in other human lung cells. We compared anchorage independence of KRAS^G12D^-AALE cells treated with or without dox by soft agar assay. Dox treated KRAS^G12D^-AALE cells (with *SMARCA4* knockdown) formed significantly fewer colonies compared to dox untreated cells (Fig. 6I). Similar results were observed when AALE-KRAS^G12D^ cells were grown under low attachment (data not shown). These results strongly suggest that SMARCA4 promotes transformative properties of KRAS^G12D^-transformed human lung cells.

### Mutational alterations in *SMARCA4* in human lung adenocarcinoma

We next evaluated the specific types of *SMARCA4* mutations observed in Lung adenocarcinoma samples. We found that missense mutations accounted for 50% of total *SMARCA4* somatic mutations (Fig. 7A). We compared this *SMARCA4* mutational profile to those of the most commonly mutated tumor suppressor genes (TSGs) in lung adenocarcinoma^22^. Unlike *SMARCA4*, the *STK11, NF1, RB1, and RBM10* genes frequently harbored truncating mutations (nonsense, frameshift, splice site mutations). *KEAP1, and TP53* were exceptions, with significantly more missense mutations. Importantly, almost 60% of missense mutations in *KEAP1* had gain of function (GOF) properties leading to elevated oncogenic NRF2 levels and altered protein interactions^23^ and *TP53* has well-established gain of function properties^24^. Due to the enrichment for missense (over nonsense) mutations in *SMARCA4*, we speculated that a subset of *SMARCA4* missense mutations might similarly have GOF or change of function properties that contribute to tumor fitness, as suggested by sequencing studies^25^. We tested this hypothesis by using the predictive algorithm, Mutation Assessor which can predict switch of function (SOF) mutations by determining if the mutated residue impacts known or predicted protein binding interfaces and hence affect functional interactions^26^. Roughly 70% of the *SMARCA4* missense mutations were predicted to have a functional impact (Supplementary Table 1). Importantly, G1232S, the most frequent *SMARCA4* mutation in lung adenocarcinoma patients^25^ (Fig. 7B, q-value=0.007), was a potential change of function mutation, indicated by a higher variant specificity score compared to variant conservation score (Supplementary Table 1). To functionally exclude the possibility that *SMARCA4*^G1232S^ encoded a loss of function mutation, we employed an auxotrophic-complementation method in a Snf2 deletion (ΔSnf2) *Sacchromyces cerevisiae* strain. ΔSnf2 is auxotrophic to galactose, which is rescued by expression of WT-Snf2. We constructed a yeast-human Snf2-S*MARCA4*^G1232S^ chimera construct as previously described^27^. We found that Snf2-WT-SMARCA4 rescued the growth of ΔSnf2 on galactose, as expected (Fig. 7C). Notably, the Snf2-S*MARCA4*^G1232S^ chimera grew better than Snf2-WT-SMARCA4 strain on galactose (Fig. 7C), eliminating the possibility of a complete loss of function effect of the S*MARCA4*^G1232S^ mutation and supporting, but not proving a selection process for a change of function coding effect.

**Figure 7.**
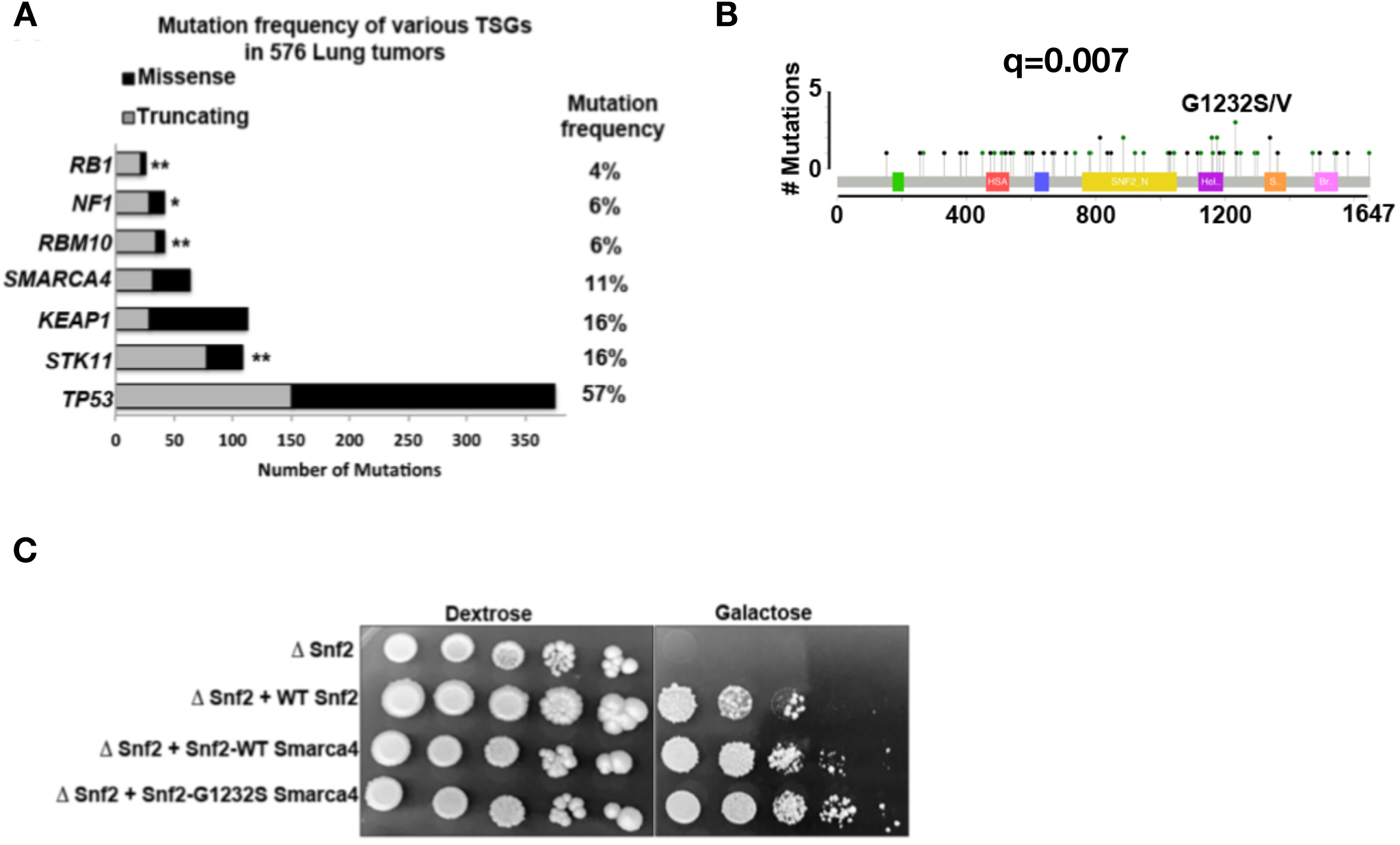
SMARCA4 mutational landscape in lung tumor. (A) Mutational frequencies of common tumor suppressor genes (TSGs) in lung ADCs. (B) SMARCA4 mutations in lung adenocarcinoma patients (C) Auxotrophic-complementation experiment in Snf2 null yeast strain with Smarca4 chimeric constructs. Ectopic expression of Snf2-1232S-Smarca4 rescues growth of Snf2 null strain on galactose

## Discussion

Identification of recurrent mutations in SWI/SNF genes in various cancers has generated considerable interest in understanding functional roles of these mutations in cancers. Understanding how these mutations contribute to cancer initiation and progression would be important in identifying vulnerabilities conferred by these mutations, to ultimately leverage this understanding to therapeutic ends. *SMARCA4* is the most commonly mutated SWI/SNF gene in lung adenocarcinoma. This cancer genomic mutational data is one of the major pieces of evidence supporting tumor suppressive roles of SMARCA4 in lung adenocarcinoma. However, we find that loss of SMARCA4 has complex and variable roles in lung adenocarcinoma progression which do not strictly conform to a typical tumor suppressor / loss of function model.

Our aim in this study was to understand how *SMARCA4* mutations influence progression of lung carcinogenesis initiated and maintained by *Kras*^*G12D*^ in GEMM models. Despite the fact that *SMARCA4* is frequently mutated in lung adenocarcinoma tumors, we find that loss of SMARCA4 limits tumor progression in mice which have undergone concomitant oncogenic *Kras* activation and *Smarca4* inactivation. Further, *Kras*^*LSL_G12D*^;*Smarca4*^*f/f*^ lung tumors were less proliferative, lower grade and showed lower levels of active KRAS signaling compared to *Kras*^*LSL_G12D*^; *Smarca4*^*+/+*^ tumors. This suggests that SMARCA4 plays a role in supporting KRAS signaling in transformed cells. We tested this by employing a two switch conditional model temporally separating *Kras*^*G12D*^ activation from *Smarca4* inactivation. We found that biallelic inactivation of *Smarca4* is selected against in *Kras*^*G12D*^-initiated lesions. A recent study used a CRISPR-based approach in the *Kras*^*G12D*^;*Trp53*^*+/+*^ (K) and *Kras*^*G12D*^;*Trp53*^*f/f*^ (KP) models to assess the loss of SMARCA4 in lung adenocarcinoma. Loss of SMARCA4 decreased tumor burden in both K and KP models. There is more heterogeneity with incomplete inactivation of target genes in CRISPR models compared to Adeno-cre activated GEMM model. Therefore, quantifying more mice (n was 4 in the study) may have provided a more robust difference in the tumor burden assay. Overall, the study concluded that SMARCA4-driven SWI/SNF functions were required continuously in KRAS^G12D^ model^28^, similar to our results but using a model with somatic editing of the allele as opposed to germline encoded Lox-P sites as we performed here.

Glaros *et al*. have previously explored the role of *SMARCA4* in carcinogen-induced lung adenomas^29^. They showed that biallelic *Smarca4* loss in untransformed cells did not increase mutagen-induced carcinoma formation. However, inactivation of *Smarca4* after carcinogen exposure lead to higher adenoma numbers when compared to *Smarca4* WT mice^29^. The difference in our observations from this study maybe due to the marked difference in mutational burden of tumors arising from ethyl carbamate (carcinogen used in the study) compared to genetic activation of *Kras*^*G12D* 30^. Further, the cell type studied differed. Glaros *et al*. uses a CCSP driven Cre recombinase, active mostly in Clara cells whereas we used Adenovirus-delivered Cre recombinase, which primarily infects type II pneumocytes, the major cell of origin for lung adenocarcinoma. We speculate that SMARCA4 may have different, and perhaps opposing, roles in these two divergent cell lineages as seen in Kras^G12D^-driven pancreatic cancer subtypes ^3,31,32^. More recently, Lissanu-Deribe et al.^33^ described the effects of loss of *Smarca4* in the setting of a Kras driver coupled with loss of *Trp53*. They observed an unprecedentedly long median survival of the comparator Kras^LSL_G12D^, *Trp53*^*flox/flox*^ (KP) animals (exceeding 16 months) making direct comparison to our studies here (which did not examine the effects of *Trp53*) or numerous previously published studies wherein KP animals always have a median survival of around than three months^34^, difficult at best.

The existence of a recurrent missense mutation at the Glycine 1232 to Serine in lung adenocarcinoma patients suggests that this mutation may have a selected for functional impact on SMARCA4. Our growth enhancing phenotype observed with the chimeric yeast Snf2 protein harboring the G1232S mutation point that this mutation is not a loss-of-function mutation. It remains to be seen how this mutation affects the functional properties of the SWI/SNF complex and if the G1232S mutation alters the DNA binding and histone sliding or eviction properties of SMARCA4.

Our findings suggest that intact SMARCA4 promotes growth of KRAS^G12D^-initiated tumors. Our findings indicate that SWI/SNF complexes play distinct roles in cancer in initiation and progression, determined in part by the temporal context of driving oncogenic event. Further, specific *SMARCA4* mutations may endow change or gain of function properties on the SWI/SNF complex. Finally, in light of our findings, SWI/SNF complexes containing the hotspot mutation in *SMARCA4* should be more thoroughly investigated as targets for therapeutic development, particularly in *KRAS*-mutant lung cancer.

## Acknowledgements

We thank Drs. Matthew Meyerson and Joshua Campbell for sharing data and Drs. Eric Snyder and Urisman Anatoly for assistance with mouse lung pathology. This work was supported by grants to E.A.C. from the American Lung Association (LCD-344261), the NCI/NIH R01CA178015, R01CA227807 and R01CA222862.

## MATERIAL AND METHODS

### Mouse studies

All experiments were approved by IACUC of the University of California, San Francisco. *Kras*^*LSL_G12D*^, *Kras*^*FSF_G12D*^, *Smarca4*^*f/f* 16^, *SPC*^*CreERT2*^ mice have been previously described^35^,^18^,^15^. Mice were bred on a mixed background. Adenovirus expressing Cre and Flippase recombinase (Viraquest, University of Iowa) were administrated into the nasal passages of mice as previously described^36^. Briefly, 2-3 months old mice were infected with 2×10^7^ or 10^8^ pfu of adenovirus expressing Cre or Flp recombinase respectively. Tamoxifen (Sigma) was administrated by intraperitoneal injection (1mg/mouse) for 5 consecutive days.

We estimated recombination efficacy achieved by *Spc*^*CreERT2*^ in mouse tumors initiated by Adeno-Flp by the mT/mG (membrane-targeted tandem dimer tomato/green fluorescent protein) reporter^37^. 45% of Adeno-Flp initiated tumors expressed GFP upon tamoxifen administration (Supplemental Fig. 1A).

### Quantification of tumor burden and tumor grading

Hematoxylin and Eosin (H&E) stained mouse lung sections were scanned using an Aperio Scanscope. Tumor burden (ratio of area of lung occupied by tumors to total lung area) as well as areas within the lung tumors showing high-grade nuclear cytologic features was quantified using ImageScope Software.

### Immunohistochemistry

Lungs were fixed overnight in Z-FIX (Anatech) at 4°C, embedded in paraffin, cut into sections and placed on slides. Following citrate-mediated antigen retrieval by boiling, slides were blocked with 5% BSA and were incubated with antibodies against SMARCA4 (H-88, Santa Cruz), Cleaved caspase 3 (Cell Signaling), Ki67 (Abcam), p-AKT (AKT pS473, Cell Signaling Technology), pERK1/2 (ERK pT202/Y204, Cell Signaling Technology), TTF-1 (Santa Cruz), overnight at 4°C in a humidified chamber. Secondary antibodies were applied for 1h at room temperature. Detection was conducted using the DAB Chromogen System (Dako).

### RNA Sequencing and Analysis

1 ug of total RNA was isolated from *Smarca4*-deficient and WT Smarca4 reconstituted mouse cell lines. Library was prepared using kappa biosystems kit. 10nM library was used for sequencing on a HiSeq 2000 instrument generating 100 base-pair paired reads. Reads were mapped to the mouse genome (mm10) using TopHat2 with default parameters. Cufflinks and Cuffdiff were used for transcript assembly and differential expression analysis respectively.

GSEA analysis was done using the GSEA software (Broad Institute). Pathway analysis was done using Metscape. Threshold fold change was set to +/- log 0.6 at 95% confidence level.

### Cell Culture

AALE cells were provided by Dr. Matthew Meyerson (Dana Farber / Harvard). All mouse lung lines were cultured in RPMI with 10% fetal bovine serum (FBS) and penicillin/streptomycin. AALE cells were cultured in SAGM media (Lonza). All other cell lines (Panc.1, SKHEP1) were maintained in DMEM with 10% FBS and penicillin/streptomycin. *Kras*^*LSL_G12D*^; *Smarca4*^*f/f*^ cell lines were generated from mouse lung tumors by excising and then digesting the tumors using 0.25% trypsin. Digested tumors were plated in RPMI media and allowed to grow for several weeks.

Single clones were then picked, propagated and probed for *Kras* recombination by PCR and SMARCA4 loss by western blot.

*SMARCA4* was re-expressed in *SMARCA4* deficient cells by transducing cells with retrovirus expressing human SMARCA4. AALE were transformed using retroviral vector pBabe-Flag-*KRAS*^*G12D*^*-*Zeo. The Zeocin selected AALE cells were then transduced with lentiviral particles containing pTRIPZ-*SMARCA4* shRNA (clone #153152_GE Dharmacon) and selected with puromycin. For western blot experiments, cells were collected 72h post doxycycline (1ug/ml) induction.

### Immunoblotting, Immunoprecipitation, and Glycerol Gradient Sedimentation

Cells were lysed using radioimmunoprecipitation (RIPA) buffer containing phosphatase and protease inhibitors. Extracts were subjected to sodium dodecyl sulfate (SDS)-polyacrylamide gel electrophoresis and then transferred to PVDF membrane. Membranes were probed for SMARCA4 (Santa Cruz), β-Actin (Santa Cruz), E-cadherin (Cell Signaling), Vimentin (Cell signaling), pAKT (AKT pS473, Cell Signaling Technology) and total-AKT (Cell Signaling) antibodies.

For immunoprecipitations, nuclear extracts were prepared by ammonium sulfate precipitation, as described ^38^. Protein precipitate was treated with DNAse for 30 minutes at 37 degrees and resuspended in IP buffer (50mM Tris pH 7.5, 150mM NaCl, 1mM EDTA, 10% glycerol, 0.5% Triton X). Lysates were incubated with anti-SMARCA4 (Abcam, ab110641) followed by addition of Dyna beads for crosslinking to Dyna beads. Beads were washed and eluted in LDS Loading Buffer (Invitrogen) for analysis by Western blotting or mass spectrometry (performed by the Salk Institute Mass Spectrometry Core). Silver stain was carried out using Invitrogen SilverQuest Silver Staining Kit. For Western blotting of SWI/SNF subunits, the following antibodies were used: SMARCA4 (Santa Cruz, sc-17796), BRM (Bethyl, A301-015A), SMARCC1 (Santa Cruz, sc-10756), SMARCB1 (Santa Cruz, sc-166165), SMARCE1 (Bethyl Labs, A300-810A), SMARCD1 (Santa Cruz, sc-135843), ACTL6A (Novus Biologicals, NB100-61628). Glycerol gradient sedimentation analysis was performed as described previously ^38^.

### Soft agar

10^5 cells (*Kras*^*G12D*^, *Smarca4*^*f/f*^ mouse lines) or 20,000 AALE cells were plated on a 0.35% low melting agarose (Lonza) placed over a 0.7% agarose layer in a 6-well plate. For AALE cells, bottom layer contained 0.5% agarose. A few drops of media were added on top every week to keep the agarose moist.

For soft agar experiments with doxycycline treatment, 100ul of media containing doxycycline (1ug/ml) was added twice/week on top of the agarose layer. Cells were incubated for two to eight weeks. Colonies were counted under a bright filed microscope for quantification.

### RNA Isolation and qPCR

Cells were homogenized using QIAshredder and total RNA was isolated using the RNA-easy kit (Qiagen). cDNA was synthesized using high capacity RNA to cDNA kit (Applied Biosystems). Q-PCR was performed with SYBR green-based gene expression assays (Roche). Primer sequences are available on request.

### Statistical Analysis

All statistical analyses were performed using GrapPad Prism, Microsoft Excel or the R statistical computing language. Statistical significance, calculated using Student’s *t*-test or Fisher’s exact test, is depicted as: * p<0.05, **p<0.01, ***p <0.001 and # >0.05

### Bioinformatics Analysis

For figure 1a, all LUAD patients (n = 575, dataset from ^21^) were classified into onconegatives and oncopositives groups. A sample was considered as oncopositive if mutations were found in EGFR, BRAF, ERBB2, MET, KRAS, MAP2K1, HRAS, NRAS, RIT1 genes and amplifications were found in MET gene. We then compared the mutation frequency in onconegative and oncopositive samples and enrichment was calculated using Fisher’s Exact test. A p-value <0.05 was considered as significant. Frame shift, missense, nonsense and splice site mutations were considered in this analysis. For figure 1b, KRAS signature (from ^13^) was applied to LUAD patients (dataset from ^39^) and lung ADCs were divided into KRAS high and low groups. SMARCA4 mutation and expression was analyzed in these 2 groups. Deleterious mutations include frameshift, nonsense and splicing mutations.

---

**Supplemental Excel 1:** Output of running Mutation Assessor algorithm on all SMARCA4 mutations found in lung ADCs.

**Supplemental Figure 1:**
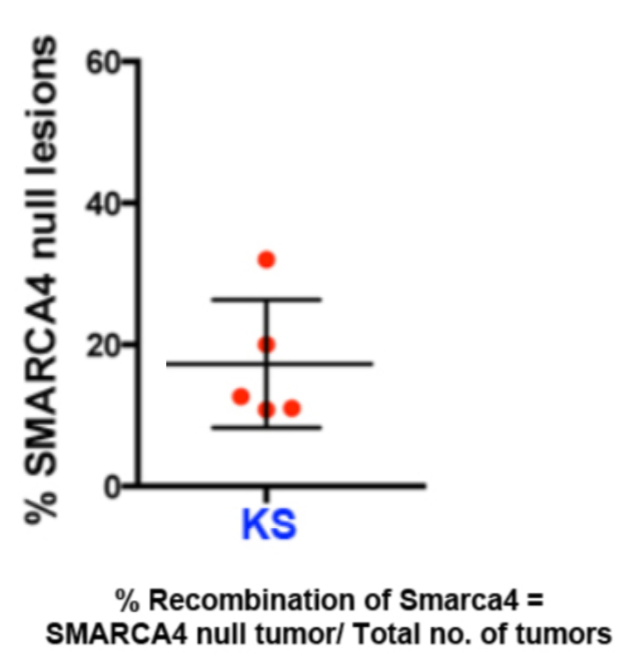
Percentage of recombination of *Smarca4* in KS mice.

